# CsoDIAq Software for Direct Infusion Shotgun Proteome Analysis (DISPA)

**DOI:** 10.1101/2021.05.12.443833

**Authors:** Caleb W. Cranney, Jesse G. Meyer

## Abstract

New mass spectrometry data collection methods require new computational tools. Direct Infusion Shotgun Proteome Analysis (DISPA) is a new paradigm for expedited mass spectrometry-based proteomics, but the original data analysis workflow was onerous. Here we introduce CsoDIAq, a user-friendly software package for the identification and quantification of peptides and proteins from DISPA data. In addition to establishing a complete and automated analysis workflow with a graphical user interface, CsoDIAq introduces algorithmic concepts to improve peptide identification speed and sensitivity. These include spectra pooling to reduce search time complexity, and a new spectrum-spectrum match score called match count and cosine (MaCC), which improves target discrimination in a target-decoy analysis. We further show that reanalysis after fragment mass tolerance correction increased the number of peptide identifications. Finally, we adapt CsoDIAq to standard LC-MS DIA, and show that it outperforms other spectrum-spectrum matching software.

---

Shotgun proteomics using liquid chromatography (LC) coupled to tandem mass spectrometry (MS/MS) is currently the leading method to identify and quantify proteome dynamics from biological samples. Two main types of mass spectrometry (MS) data acquisition exist, namely data-dependent analysis (DDA)^1–3^ and data-independent analysis (DIA)^4–6^. As the name implies, the scan sequence in DDA depends on the data; even if the same sample is analyzed twice, the scans collected in each analysis are unique. In each DDA scan cycle, the MS surveys *m/z* values that may represent peptides in a precursor scan, followed by isolation and fragmentation of those *m/z* regions in tandem mass spectrometry (MS/MS) scans. In contrast, DIA scans are the same in each analysis. DIA fragments all masses within a predefined set of *m/z* ranges, usually spanning the range of useful peptide masses from approximately 400-1,000 *m/z*. DIA scans therefore usually result in chimeric spectra representing the combined MS/MS spectra of multiple peptides. DIA has grown significantly in popularity since its conception, as DIA data allows for deeper and more consistent peptide quantification than DDA in the absence of peptide fractionation^7^. However, methods for DIA data analysis are still maturing, and continued advancements are required to maximize the value extracted from DIA. Further, the continued development of new DIA data collection methods requires specialized new tools.

Several methodologies exist for identifying peptides from DIA MS data, including EncyclopeDIA^8^, PECAN^9^, Spectronaut^10^, DIA-Umpire^11^, DIA-NN^12^, Thesaurus^13^, OpenSWATH^14^, Skyline^15^, mProphet^16^, LFQbench^17^, and PIQED^18^. Recent advances in machine learning have opened up the possibility of *de novo* sequencing^19^, or matching to predicted MS/MS spectra, such as Prosit^20^, DeepMass^21^, and DeepDIA^22^; however, many DIA data analysis methods require scoring the presence of peptides by comparing to spectra previously identified by DDA. Because nearly all proteomics DIA relies on LC, this is often achieved by assigning possible peptides a score based on the co-elution of peptide fragment ion signals over time. Retention time plays an important role in limiting the search for peptide fragment signals^23^. True and false peptide matches are segregated using the target-decoy strategy to estimate false discovery rate (FDR)^24^. A different strategy that only considers each spectrum without need for LC uses the projected spectrum concept^25^. MSPLIT-DIA^26^ identifies peptides from complex, chimeric DIA spectra by comparing the shape of only those fragment ion intensities that are found within some mass tolerance of library spectra fragments.

Nearly all proteomics experiments rely on LC to separate peptides before ionization and MS analysis. The field of proteomics is experiencing a trend toward shorter LC gradients^27,28^. The logical extreme is to remove LC entirely; we recently introduced a new paradigm that enables fast proteomics called direct infusion shotgun proteome analysis (DISPA), which does not use LC separation and instead relies on additional gas-phase fractionation by ion mobility^29^. Because direct infusion data lacks co-elution of peptide fragments over time, the original report of DISPA relied on projected cosine scoring with MSPLIT-DIA for peptide and protein identification. However, because MSPLIT-DIA was not customized to DISPA data and does not natively identify proteins, multiple custom python and R scripts were required to enable FDR calculation, protein identification and quantification. Overall, the original data analysis process was inaccessible and could deter future use of DISPA, despite its potential to enhance our study of the proteome.

Here, we describe CsoDIAq (Cosine Similarity Optimization for DIA qualitative and quantitative analysis), a python software package designed to enhance usability and sensitivity of the projected spectrum concept originally utilized by MSPLIT-DIA. CsoDIAq introduces several algorithmic advances, including pooling spectra peaks for reduced time complexity and a new spectra-spectra scoring function that improves discrimination of target and decoy peptides. Combined with a Graphic User Interface (GUI), CsoDIAq is both effective and user friendly, and can analyze DIA data from DISPA and LC-MS. We show that CsoDIAq identified 23.3% more peptides than MSPLIT-DIA when applied to DISPA data, and 36.0% more peptides from standard LC-MS DIA data (FDR < 1%).

## METHODS

### Data and Formats

CsoDIAq reads raw mass spectrometry data in mzXML format, and spectral libraries created with SpectraST^30^ in TraML tsv format are preferred. However, we also support mgf libraries created with MPLIT-DIA^26^, or the pan-human library^31^. CsoDIAq requires .csv to be comma-delimited and .tsv files to be tab-delimited. Because downloads of the pan-human library included .csv files that are tab-delimited, some libraries may need to be converted to the appropriate format. Spectral libraries were generated with multiple settings and the best library creation settings were: no fragments corresponding to loss of water/ammonia, only fragments from 400-2000 *m/z* within a 0.2 *m/z* tolerance of the predicted mass (the initial TraML library was built from low-resolution ion trap MS2 data).

DISPA data used to develop CsoDIAq was from the original publication^29^. It should be noted that libraries with a greater excess of nonexistent peptides will identify fewer peptides^32^. In our case, the TraML library used in below analyses has fewer peptides and generally outperforms the mgf library. Spectral libraries and all code and data to reproduce the figures are posted to a new repository on zenodo.org (10.5281/zenodo.4750187). LC-MS data was downloaded from massive^33^.

### Spectra Pooling

CsoDIAq introduces a library-query peak comparison method dubbed “spectra pooling” that reduces the time complexity by an exponential factor. Four variables primarily impact the speed of the algorithm in any given *m/z* window of a DIA analysis, namely the number of library spectra corresponding to that window (nLS); the total number of fragment ion peaks in nLS library spectra (pLS); the number of query spectra (nQS); and the total number of fragment ion peaks in nQS query spectra (pQS). MSPLIT-DIA^26^ iteratively compares each library spectrum to each query spectrum, presuming the precursor mass of the peptide represented by the library spectrum falls within the *m/z* window captured by the query spectrum. If the above variables are assigned the letter values of nLS, pLS, nQS, and pQS, respectively, the time complexity of this method would be:

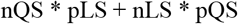

Variation in these factors significantly impacts the length of time required to complete the algorithm. In big O notation, the above equation results in a time complexity of O(n*m).

Spectra pooling reduces unnecessary repetition in peak comparison, significantly improving speed at no cost to accuracy. MSPLIT-DIA separately compares a query spectrum to each relevant library spectrum, therefore referencing the same peak from one spectrum type once for each other spectrum with a precursor *m/z* within a given *m/z* query window. Spectra pooling instead assigns each fragment ion a spectrum tag in addition to its inherent mass and intensity values, which allows consolidation or pooling of multiple spectra into a single spectrum for comparison. Matches to fragments in the pooled spectra can be separated after matching using their spectrum tag to compute the separate match scores. Thus, by comparing a pooled query spectrum to a pooled library spectrum, any peak would only ever be referenced once. This exponentially reduces the time complexity of the above conventional approach to:

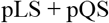

In big O notation, this results in a new time complexity of O(n+m).

DISPA scouting experiments iterate over the same *m/z* query window at least once for every FAIMS compensation voltage setting. In terms of the above equation, nQS is generally equal to the number of compensation voltage settings run in the experiment. The dataset used as the benchmark iterated over the same *m/z* query windows twelve times for a scouting experiment, twice each for six compensation voltage settings.

Two additional versions of the algorithm, one with only library spectra pooling and one with no pooling, were created to graphically illustrate the impact of spectra pooling on time complexity. Only pooling one spectrum type enables graphical comparison of performance between pooling and non-pooling on spectra from the other type. We pooled library spectra as opposed to query spectra for graphical representation because the number of library spectra (generally measured in thousands) often far exceeds the number of query spectra (approximately six) in a given *m/z* query window for DISPA data, and thus will more fully demonstrate the dramatic reduction in time complexity. Both versions of the algorithm, spectra pooling and non-pooling, were based on copies of the main algorithm, which was created under the assumption that pooling would occur. We did not optimize the non-pooling algorithm contrary to this expectation, which may cause additional time lag. However, the overall reduction in time complexity remains as above described.

Query spectra are grouped by precursor *m/z* and window width for pooling. By default, CsoDIAq pools all grouped query spectra, but users can indicate a maximum number of spectra to pool to reduce memory use.

### Scoring Method

CsoDIAq employs a new scoring function for spectra-spectra matching (SSM) that improves segregation of target and decoy peptide distributions to optimize the number of peptide hits below a standard False Discovery Rate (FDR) cutoff. CsoDIAq first takes the square root of fragment ion peak intensities in the spectral library and experimental spectra to normalize the contributions of fragment ion intensities^34–36^. Next, for each experimental spectrum, the fragment ions are compared with the pooled library spectra of all possible matches. Like MSPLIT-DIA, CsoDIAq uses the ‘projected spectrum’ concept; only experimental fragment ions found within a defined mass tolerance of fragment ions in the library spectra are used to compute the score. Fragment comparisons are done using parts per million (PPM) rather than absolute *m/z* differences. All matched fragment ions are recorded and then used to compute a SSM score for all possible peptides in the pooled library. CsoDIAq calculates a SSM score by multiplying the fifth root of the number of matched fragments by the cosine score. Because of the importance and impact of peak matches on the SSM score, CsoDIAq imposes a minimum of three fragment ion matches to the library spectrum with no maximum.

The number of matches between a library and query spectra plays a significant role in these calculations, and as such noise in a library spectrum can strongly skew the MaCC score. This is primarily a concern for MGF libraries, as the TraML format already filters for fragment mz values expected for a given peptide. As such, all libraries are pre-processed to only include the top ten most intense peaks. For the same reasons, CsoDIAq will only function with centroided experimental data.

### Fragment Mass Error Correction Process

CsoDIAq also employs a dual search strategy for fragment ion mass correction. When comparing library peaks with query peaks, *m/z* values for true corresponding fragments are not expected to precisely match. In addition to a general margin of error in the query spectrum resulting from the natural variance of mass spectrometers, drift in mass calibration can result in a systematic mass value offset. To adjust for this, CsoDIAq runs an initial, uncorrected analysis of the data using a generic offset of 0 PPM and a default, user-adjustable tolerance of 30 PPM. These numbers were based on previous experimentation that suggested an overall window of 30 PPM around 0 PPM would capture the true offset in addition to sufficient data to calculate an optimized tolerance. After identifying peptides of interest using the previously described scoring method, csoDIAq determines a new offset and tolerance from the PPM differences for those hits. By default, the offset and tolerance is customized to the PPM distribution. The offset is the highest bin value of the given histogram and tolerance is the furthest bin from the offset with values approximately equal to the surrounding noise. Users can alternatively elect to use the median and standard deviation of their choice for the offset and tolerance, respectively. CsoDIAq then excludes all peak matches outside of the chosen PPM offset and tolerance, resulting in a corrected analysis that outperforms the uncorrected analysis in the number of unique identifications.

For reference, the MSPLIT-DIA has a minimum allowance of ten matching peaks and a minimum cosine score of 0.7. Results were sorted by cosine score to calculate the FDR of each SSM for comparison with CsoDIAq output. Notably, all SSMs from the MSPLIT-DIA consistently had a lower FDR than 0.01, leading to acceptance of all SSMs.

### Peptide and Protein Identification

CsoDIAq produces three output files for each mass spectrometry file that report spectra, peptides and proteins filtered to <1% FDR. In each case, CsoDIAq sorts peptide identifications by the above-described score, calculates the FDR for each identification using a modification of the target-decoy approach where FDR at score *S* = # decoys/ # targets, and removes SSMs below a 0.01 FDR threshold. The spectra report is returned without filtering by unique peptides. The peptide FDR calculations only use the highest-scoring instance among all SSMs for each peptide. CsoDIAq uses the IDPicker algorithm^37^ to identify protein groups from the list of discovered peptides and adds them as an additional column in the output. Protein groups from the TraML spectral library are used for protein inference rather than matching peptides back to protein entries in a FASTA file. Our implementation of the IDPicker algorithm preferentially identifies proteins with a higher number of peptide connections after the peptide reduction step. When there is a tie, the algorithm instead uses the original number of peptide connections per protein. The protein FDR calculation for proteins uses the highest-scoring peptide as the protein group score, though all peptides connected to those proteins in the peptide FDR output are re -included in the protein report for reference.

### Protein Quantitation

Accurate protein quantitation requires a second DIA data collection that targets *m/z* and Compensation Voltage (CV) values corresponding to the best peptide target from identified proteins. CsoDIAq uses two criteria to choose representative peptides identified for each protein. First, peptides not unique to a given protein are eliminated from consideration. Next, CsoDIAq sorts the peptides from each protein by summed fragment ion intensity of matched fragment ions. Finally, the software allows the user to input their desired maximum number of representative peptides from each protein, starting with the highest ion count.

The targeted quantitative DISPA re-analysis currently requires that samples be prepared using Stable Isotope Labeling by Amino acids in Cell culture (SILAC)^38^, specifically using both ^13^C_6_, ^15^N_2_ lysine and ^13^C_6_, ^15^N_4_ arginine. CsoDIAq first prepares library spectra specific to the y-ions of the targeted peptides and their heavy isotopes. CsoDIAq uses a default initial tolerance of 30ppm before optionally applying the same mass correction algorithm discussed earlier to identify an offset and tolerance specific to the DISPA run. After identifying matched peaks (default: at least one of the top three most intense peaks), CsoDIAq calculates the SILAC ratio for each peptide based on the identified peaks (default: median ratio value).

The user can input (1) the standard deviation used to determine the tolerance of the correction process, (2) the minimum number of matches required to calculate SILAC ratios for the peptide, and (3) the mode of ratio selection.

Note that the file for the targeted re-analysis will not have all the leading proteins from the protein FDR file. This is because the decoys will be removed, and because some protein groups identified by the IDPicker algorithm won’t have unique peptide targets to use.

### Comparison with MSPLIT-DIA

For comparing the output of MSPLIT-DIA with CsoDIAq we generated an MGF library using the original data pipeline from skyline .blib converted to .ms2 and then .mgf. Peptides in the library were stripped of modifications for protein identification from a FASTA file after initial library generation. The script for adding proteins to an MGF file is included in the CsoDIAq package at the command line. Both program settings included an initial tolerance of 10 PPM. MSPLIT-DIA was set to 2 *m/z* precursor window width for the DISPA data and 9 *m/z* window width for the LC-MS DIA data, both of which used a setting of 0 scans per cycle. CsoDIAq takes the window width from the mass spectrometry data file. Aside from the initial tolerance, all default settings were used for CsoDIAq output. MSPLIT-DIA output was processed using the same FDR calculation algorithms used by CsoDIAq at both the peptide and protein level. The MSPLIT-DIA output column name “Peptide” was altered to “peptide” for this process, and the output was sorted by the cosine score rather than the MaCC score for FDR calculation.

### Usability

CsoDIAq was written to be used from the command line through the pip installation package. All help text and flag descriptions can be viewed with the “--help” flag, as is standard for programs triggered from the command line. The CsoDIAq command line returns an error for improper inputs.

In addition to command line operations, the CsoDIAq software package includes a graphical user interface (GUI) implemented with the package PyQt5^39^. Ultimately, the GUI only serves as a shell for command line prompts and flags. From the GUI, invalid inputs highlight the offending section title in red, whereas the command line throws an error. A text window included in the GUI indicaties progress through the program and highlights errors should they arise.

### Code and Availability

CsoDIAq is written using the Python computer programming language. The program utilizes several software packages to enhance time and memory efficiency. For example, the numba package^40^ allows the program to run with time efficiency approaching those written with coding languages such as C++ and Java and use of the numpy^41^ package drastically reduces memory use. Additional packages used include matplotlib^42^, pyteomics^43^, pandas^44^, biopython^45^, PyQt5^39^, and lxml^46^. CsoDIAq is available on GitHub (https://github.com/CCranney/CsoDIAq).

## RESULTS

### Overview

DISPA has emerged as a promising method for fast peptide and protein identification and quantitation. However, the original pipeline lacked unified computational support (**Supplemental Figure 1**). As shown in **Figure 1** and as expounded in the original paper, DISPA data analysis requires: (1) spectra-spectra match scoring, (2) FDR filtering to remove false positives, (3) protein inference, and (4) peptide quantitation via light-to-heavy isotope ratio comparison. In the original implementation, MSPLIT-DIA^26^ was used to score peptides from DISPA data according to the projected spectrum concept^25^. However, an integrated solution including the above-mentioned workflow steps was not developed in the initial publication. To address this limitation, here we introduce CsoDIAq (Cosine Similarity Optimization for DIA qualitative and quantitative analysis), a software package that efficiently consolidates the analytical pipelines needed for DISPA data analysis (**Figure 1**). In addition to supporting the steps that previously required custom code, CsoDIAq optimizes peptide identification by introducing a new scoring function based on a combination of projected cosine and the number of matched fragment ions. Further, we introduce a 2-stage search strategy including fragment mass tolerance correction. The combination of these new features, along with the ability to search against an improved TraML library, nearly doubles the number of unique peptide identifications at <1% FDR compared to the original publication using the same data.

**Figure 1:**
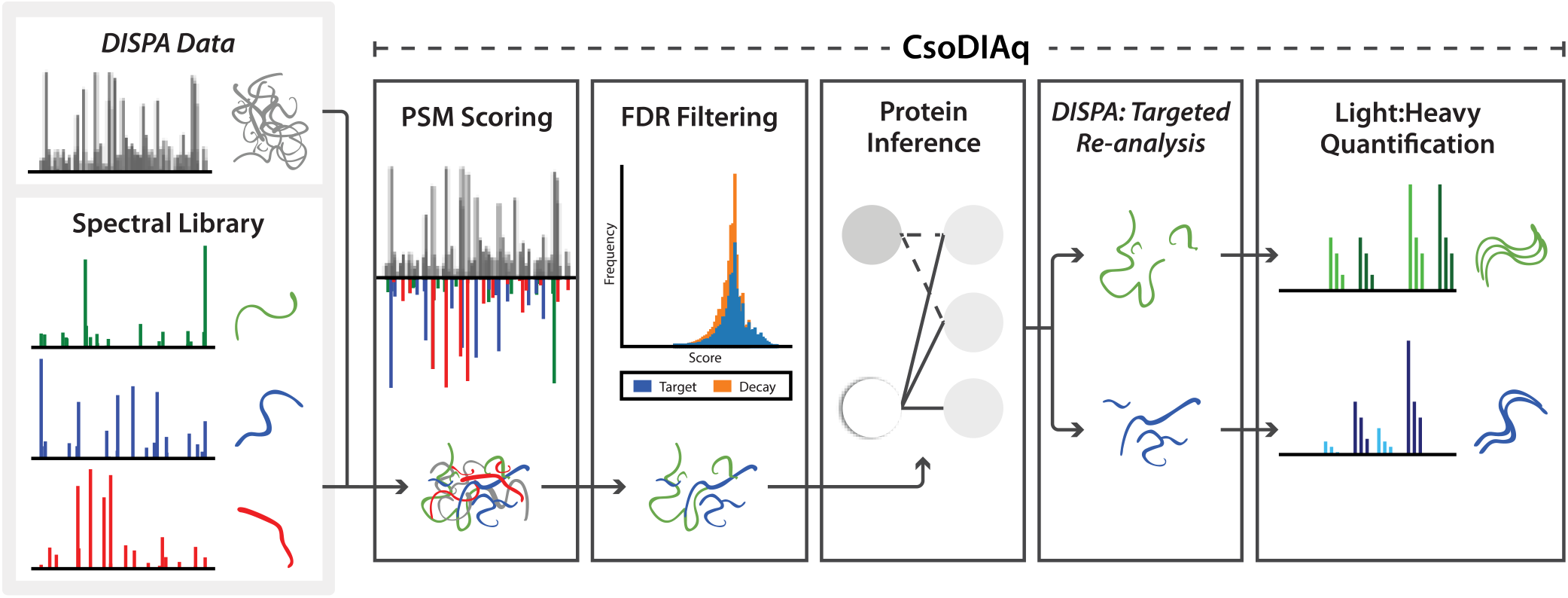
Overview of CsoDIAq software package.

### Spectra Pooling

The use of spectra pooling to compare library and query spectra significantly improved the time performance of the algorithm. Rather than iteratively comparing multiple library spectra to multiple query spectra, spectra pooling tags each peak with a spectrum-specific identifier to enable library spectra “pooling”. Key to this strategy is subsequent fragment ion match separation for scoring. By pooling all relevant spectra prior to peak comparison, CsoDIAq only ever iterates over each fragment ion peak a single time, which reduces the time complexity from O(m*n) to O(m+n) (**Figure 2A**). CsoDIAq analysis was performed with and without pooling to illustrate this effect. **Figure 2B** shows how the time required to compare a single experimental spectrum increases with the number of query fragment ions and the number of potential library spectra matches. The number of fragment ion peaks in the query spectra shows positive linear correlation with time, the slope of which decreases as the number of library spectra increases **(Figure 2C)**. In contrast, the time to analyze one experimental spectrum requires roughly a static amount of time independent of the potential library spectra matches and queried experimental fragment ions (shown in [red] near the y-axis). Overall, the pooling strategy is nearly two orders of magnitude faster than without pooling (**Figure 2D**).

**Figure 2:**
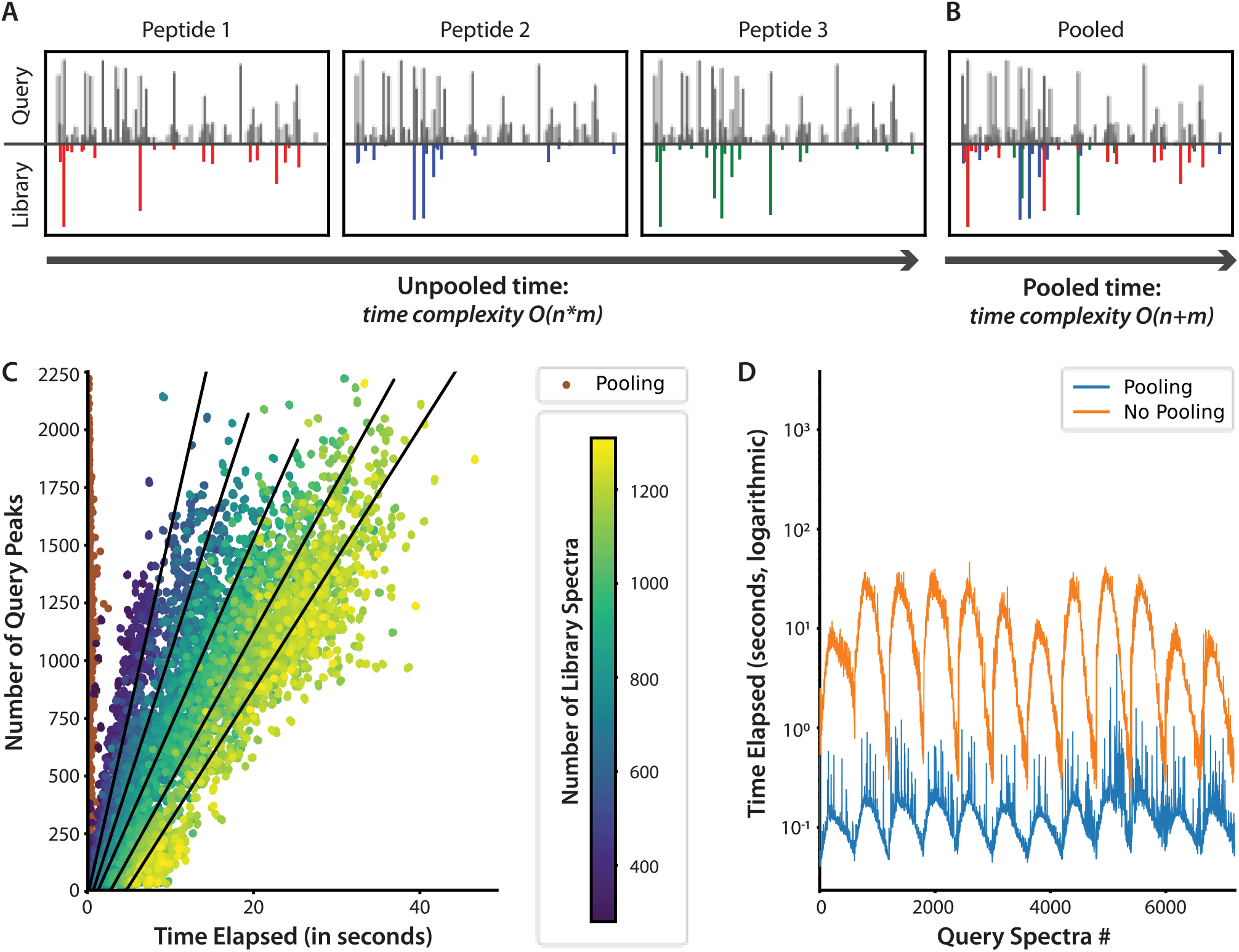
Spectra pooling concept and performance. **A**, Concept figure illustrating DISPA analysis without pooling. With only one query spectrum (black) and three library spectra (red, blue, and green), the algorithm completes three iterations over all query peaks, once for each library spectrum. This indicates an O(n*m) time complexity. **B**, Concept figure illustrating DISPA analysis with pooling. Using the same spectra examples in 2A, the number of spectra no longer multiplies the number of iterations over peaks. Instead, the time complexity relies on a single iteration of all query peaks and all library peaks, indicating an O(n+m) complexity. **C**, scatterplot illustrating the effect of the number of queried fragment ion peaks and number of potential library spectrum matches on algorithm speed per single query spectra analysis. Lines indicate the slope of a subset of the data sorted by the number of library spectra in equal proportions. **D**, line graph comparing performance of pooling vs. non-pooling algorithms in regards to time. Y-axis illustrated on a logarithmic scale for visual purposes. Data shown represents time before further optimization with the numba software package.

### Peptide Spectrum Match Scoring

CsoDIAq introduces two methods that, when combined, consistently improve discrimination of target and decoy SSMs to increase peptide identifications at an FDR threshold of 1%: (1) a scoring method unique to CsoDIAq and (2) fragment ion mass corrected re-analysis.

Two variables strongly impacted the differentiation of target and decoy SSMs: (1) the number of fragment ion matches between the library and query spectra, and (2) the projected cosine similarity score. Projected cosine score was a strong indicator for identifying targets, and a higher number of fragment ion matches generally led to projected cosine scores concentrated near the optimal value of 1 (**Figure 3A**). Like MSPLIT-DIA, one tactic for separating target peptides from decoys involves filtering SSMs for a minimum number of matched fragment ions and sorting by projected cosine score. One naive approach to optimizing this tactic is to calculate the number of unique peptide identifications with an FDR cutoff of 0.01 after filtering at every possible minimum number of fragment ion matches (**Figure 3B**). A strategy introduced here is to combine the number of fragment ion matches with the cosine score, dubbed Match Count and Cosine (MaCC) score. To better equalize the contribution of the matched fragments and the cosine similarity, MaCC uses the fifth root of the matched fragment ions count multiplied by the cosine score. The MaCC score therefore results in a non-rectangular segregation of accepted and rejected hits in the space of cosine similarity versus number of matched fragments (**Figure 3C**).

**Figure 3:**
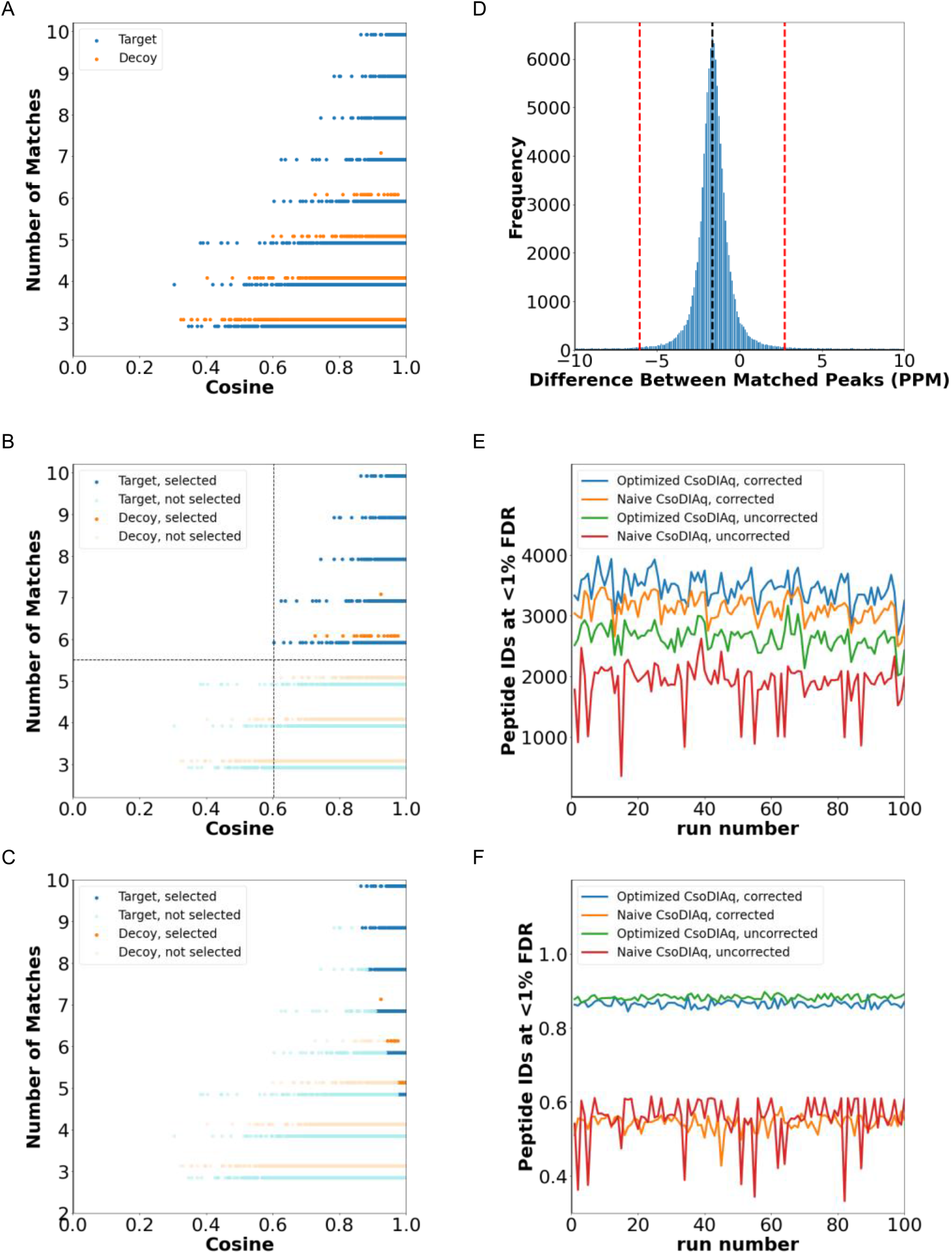
Optimized CsoDIAq scoring mechanism. **A**, scatter plot between the number of fragment matches and cosine score of all PSMs in a single run. Target and decoy PSMs visually separated as blue and orange, respectively. **B**, recreation of figure 3A indicating PSMs above an FDR rate of 0.01 as sorted using the initial scoring approach. The initial approach iteratively filters different match numbers, sorts by cosine score, and calculates the number of PSMs above the FDR rate, ultimately determining the optimal fragment ion match number to filter by. **C**, recreation of figure 3A indicating PSMs above an FDR rate of 0.01 as sorted using the optimized scoring approach, the Match Count and Cosine (MaCC) score. The optimized approach generates a new score as the fifth root of the match number multiplied by the cosine score, sorts by this new score, and calculates the number of PSMs below the FDR rate using the target decoy strategy. **D**, histogram illustrating the spread of fragment ion mass differences in PPM from all fragments corresponding to PSMs identified above the FDR cutoff rate from the first, uncorrected analysis. The black dashed line gives the mean and the dotted red lines give the second standard deviation. **E**, Line graph comparing the performance of various algorithms with respect to the number of peptide identifications over 100 DISPA runs of the same sample. A minimum number of 6 matches was applied to the naive approach to optimize the output. Like the MaCC score use, a maximum of 10 peaks per library reference was used. **F**, Line graph comparing the performance of various algorithms with respect to the lowest cosine score over 100 DISPA runs of the same sample.

After determining all SSMs with FDR<0.01 as determined by the MaCC, CsoDIAq conceptually runs a second, corrected spectrum-spectrum matching that further improved the number of identifications produced by csoDIAq. To speed up the implementation, this is not a re-comparison but rather re-filtering the matched fragments based on recorded mass errors from the initial search. A histogram of true minus predicted fragment ion mass differences in PPM of all fragmentation ion peak matches from the identified SSMs showed that mass difference was normally distributed, and that optimization of the initial range could exclude outlier fragment matches (**Figure 3D**). After refiltering, the fragment ions using the optimized fragment mass tolerance, csoDIAq’s MaCC score further excluded decoys resulting in the consistent identification of more unique peptides than all other methods (**Figure 3E**, blue line, “Optimized CsoDIAq, corrected”). Fragment mass correction also enhanced and stabilized the naive approach (**Figure 3E**, orange line, “Naive CsoDIAq, corrected”).

In addition to obtaining more peptide identifications, the combination of MaCC score and fragment ion mass correction consistently resulted in a minimum projected cosine score higher than that obtained using the naive approach (**Figure 3F**).

### Comparison with MSPLIT-DIA

Peptide and protein identifications from CsoDIAq were benchmarked against MSPLIT-DIA using the same MGF library for both DISPA and LC-MS DIA data^33^. For DISPA data, CsoDIAq identified 23.3% and 5.6% more peptides and protein groups, respectively (**Figure 4A** and **Figure 4B**); for LC-MS DIA data CsoDIAq identified 36.0% and 24.6% more peptides and protein groups, respectively (**Figure 4C** and **Figure 4D**). Tables of identified proteins and peptides are in the supplement (**Supplement Tables 1-4**). CsoDIAq produced more identifications even across the 100 replicates dataset used in **Figure 3E (Supplemental Figures 2 and 3)**.

**Figure 4:**
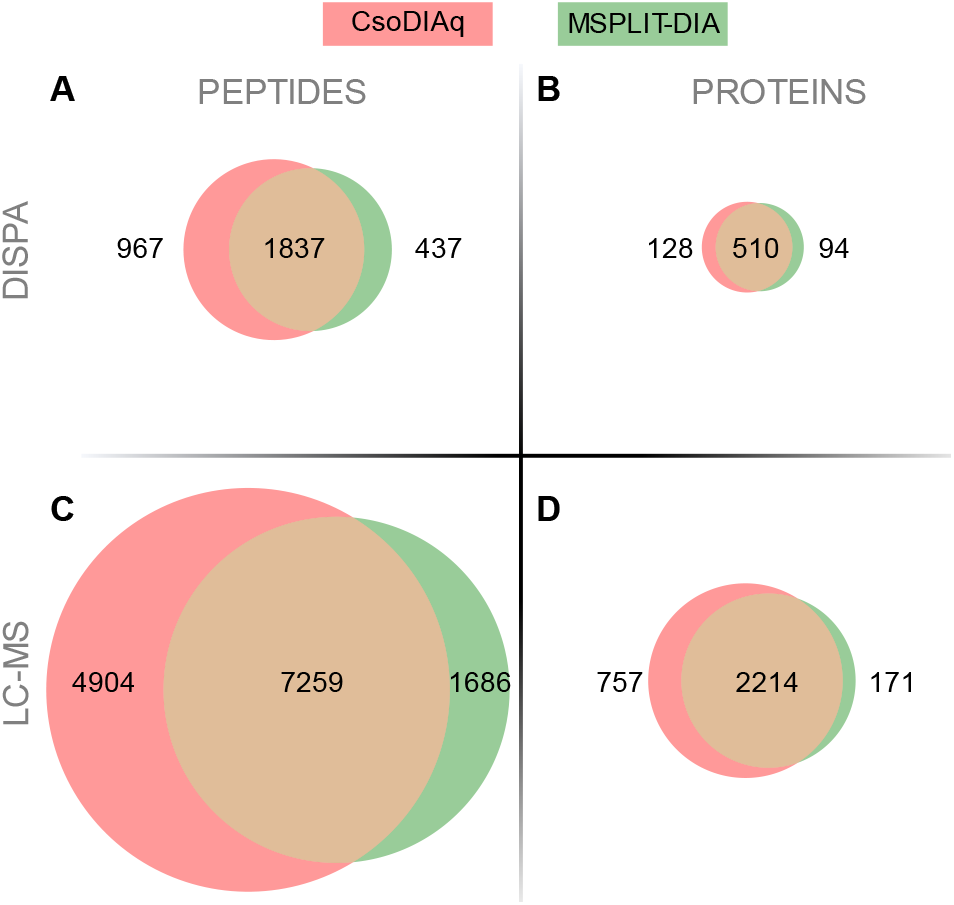
Comparing of CsoDIAq and MSPLIT-DIA performance. All values were identified above an FDR threshold of 0.01. **A**, venn diagram comparing unique peptides identified by CsoDIAq and MSPLIT-DIA using DISPA data. **B**, venn diagram comparing unique protein groups identified by CsoDIAq and MSPLIT-DIA using DISPA data. **C**, venn diagram comparing unique peptides identified by CsoDIAq and MSPLIT-DIA using LC-MS DIA data. **D**, venn diagram comparing unique protein groups identified by CsoDIAq and MSPLIT-DIA using LC-MS DIA data.

We also recorded and compared the time required to run MSPLIT-DIA and CsoDIAq for the above analyses. MSPLIT-DIA ran for 0:03:32 and 2:13:37 for DISPA and LC-MS datasets, respectively. In comparison, CsoDIAq with correction ran for a substantial improvement of 0:03:16 and 0:51:23. Because the correction process is optional, users can decrease run time at the expense of the number of targets identified above a 0.01% FDR rate. Uncorrected, CsoDIAq ran at 0:03:08 and 0:41:13.

### Protein Quantitation

CsoDIAq additionally enables peptide and protein quantification through analysis of SILAC data. We investigated the impact of three settings on the final output to determine the best defaults to use in the program: (1) the standard deviation used to determine the tolerance of the correction process, (2) the minimum number of matches required to calculate SILAC ratios for the peptide, and (3) the mode of ratio selection. We evaluated the impact of a range of these settings on the quantity and accuracy of peptide identification and quantification. Quantification data from the original DISPA publication^29^ was derived from peptide mixtures with light:heavy ratios of 1:8, 1:4, 1:2, 1:1, 2:1, 4:1, and 8:1, allowing for relative quantification of samples with known ratios. The second logarithmic of these ratios are −3, −2, −1, 0, 1, 2, and 3, respectively. With that background, the three metrics used to evaluate the CsoDIAq output were (1) the average standard deviation around the mean second logarithmic ratio across each sample, (2) the number of peptides quantified, and (3) the slope of mean second logarithmic ratios across groups.

Uniform manifold approximation and projection (UMAP)^47^ was used to reduce the multi-dimensional results into two dimensions for visualization, and the points were colored by the metrics of interest (**Figures 5A-C**). The UMAP analysis shows four groups of parameter settings; the top right has the lowest standard deviation (**Figure 5A**) at the expense of total peptides quantified (**Figure 5B**). The bottom-left group of parameters produced a slow closest to 1 (**Figure 5C**) at the expense of a high standard deviation (Figure 5A). The top-left group had good standard deviation (**Figure 5A**) and a high number of quantified peptides, but the slope of log2 ratios suffered (**Figure 5C**). The fourth group in the middle was an ideal balance between all the metrics of interest (shown as red points); these parameters were screened for those that produced (1) an average standard deviation less than 1, (2) quantified peptides greater than 80% of the maximum number of peptides identified, and (3) a slope within 0.05 of 1. This balanced quantification result was achieved using the customized PPM distribution method described as the default for identification searches, a minimum of one of the three most intense light or heavy fragment ions and using the median ratio of matched light and heavy peaks as the representative ratio (**Figures 5A-C**).

**Figure 5:**
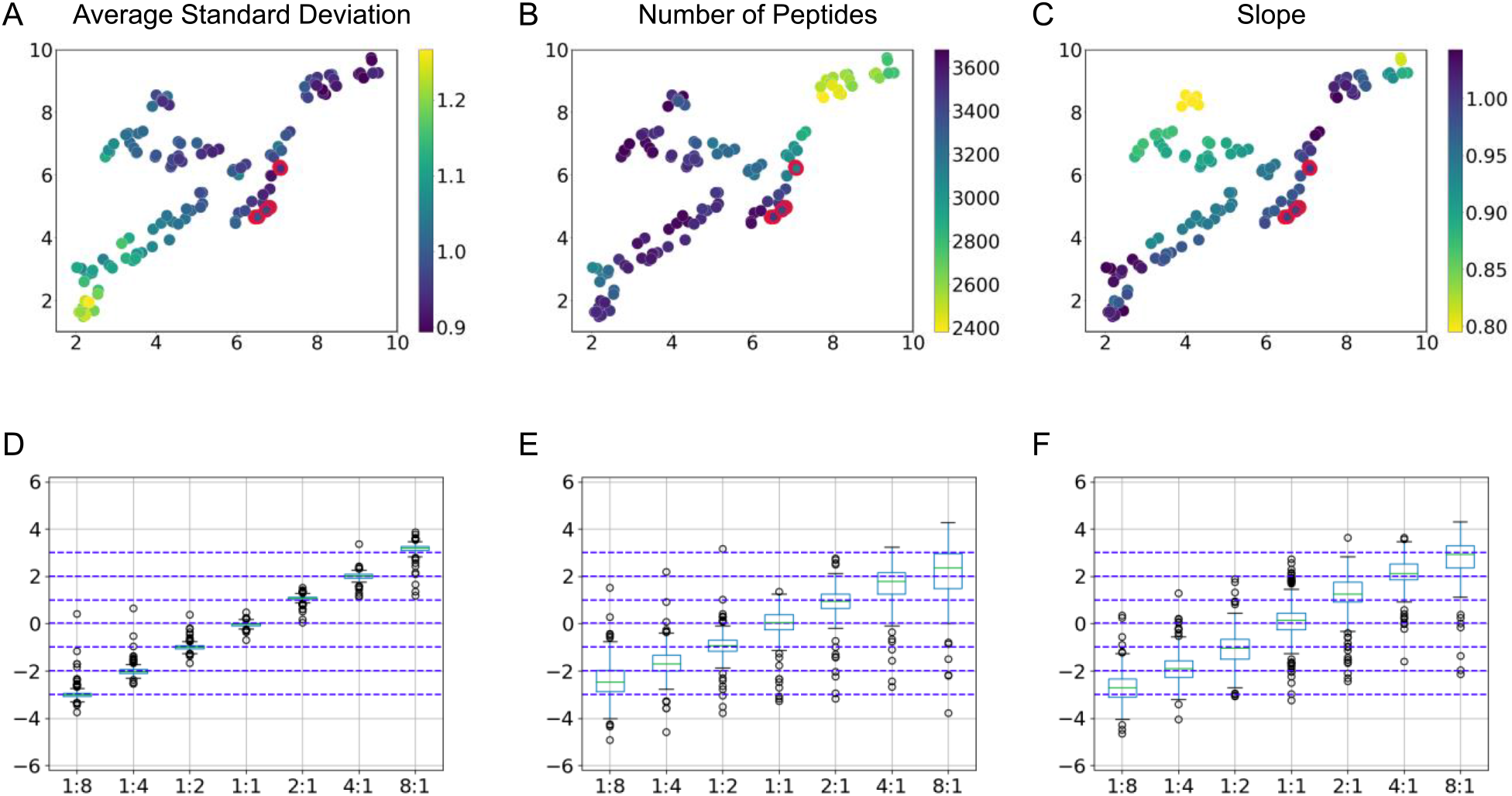
Optimizing performance of the CsoDIAq quantification algorithm. Figures A-C show UMAP that identify the best settings for use of the quantification algorithm by illustrating their impact on three most desirable outputs. Each dot represents a different mixture of settings, and red highlights indicate settings that maximize the three outputs. **A**, the impact of CsoDIAq settings on the average standard deviation around the expected second logarithmic output. **B**, the impact of CsoDIAq settings on the number of peptides quantified. **C**, the impact of CsoDIAq settings on the average expected second logarithmic output for each condition, summarized as the slope. Figures D-F are boxplots illustrating the performance of the CsoDIAq quantification algorithm in relation to other methods. They show the relative quantification of peptides for each of the mixture ratios provided. Only the 226 peptides matched across every condition were used for more direct comparison. **D**, the relative quantification as determined by standard LC-MS DDA methods. **E**, the relative quantification as determined through MSPLIT in the original DISPA publication. *F*, the relative quantification as determined through CsoDIAq.

Using output generated from the optimal settings, we next compared the results with those generated from other algorithms. Because the original DISPA publication conducted a similar analysis, we obtained the same dataset and performed a re-analysis for comparison with CsoDIAq output. After isolating to peptides identified across all analyses, we compared the relative abundance in each sample. The ratios from LC-MS match the predicted values best (**Figure 5D**) compared to the original DISPA analysis (**Figure 5E**), and our optimized algorithm showed slightly less ratio compression apparent at the extreme ratios (**Figures 5F**).

### Usability

Recognizing that isolating CsoDIAq usability purely to the command line could alienate researchers unaccustomed to such tools, we implemented a Graphic User Interface (GUI) as an aid. The GUI doesn’t add any new functionality to CsoDIAq, but serves as a shell for command line prompts to enhance usability. There are two tabs on the GUI, one for peptide/protein identification (**Figure 6A**) and one for SILAC quantification (**Figure 6B**). Both tabs have entries for DIA data file path (first DIA run for identification, second DIA run for quantification), library file path, outfile directory path, all of which are required fields. Additionally, the quantification tab requires the path to the identification tab output to match results with the prior identification. For runtime settings, defaults are used if no user input is provided and are therefore not required fields. Both tabs have settings for initial mass tolerance in ppm and correction settings, including enabling the second mass corrected analysis, standard deviation tolerance used for the correction, and whether a histogram will be generated demonstrating the PPM values specified in the correction. Identification settings also include if protein inference should be enabled, and if so, how many target peptides per protein should be included in the output target lists for quantitative analysis. There is also a setting to instigate a maximum number of query spectra that can be pooled at any time, as particularly large DIA data files can be memory intensive to analyze. The last identification setting is a checkbox that enables the files generated for targeted re-analysis to include targets for heavy peptides (includes heavy lysine and arginine). Quantification settings include an entry for the maximum number of library peaks per library spectra and a minimum number of peak matches required for identification and quantification, as excess peak matches decrease quantitative accuracy. Because each fragment match between library and query spectra can be used to determine a ratio that represents the change in quantity between conditions, a setting to choose between the mean or median of matched peak ratios is included as well. In all cases, invalid inputs are highlighted red after clicking the “Execute” button while valid inputs are highlighted green. Conditions required for each field can be identified by hovering over the highlighted text field in question.

**Figure 6:**
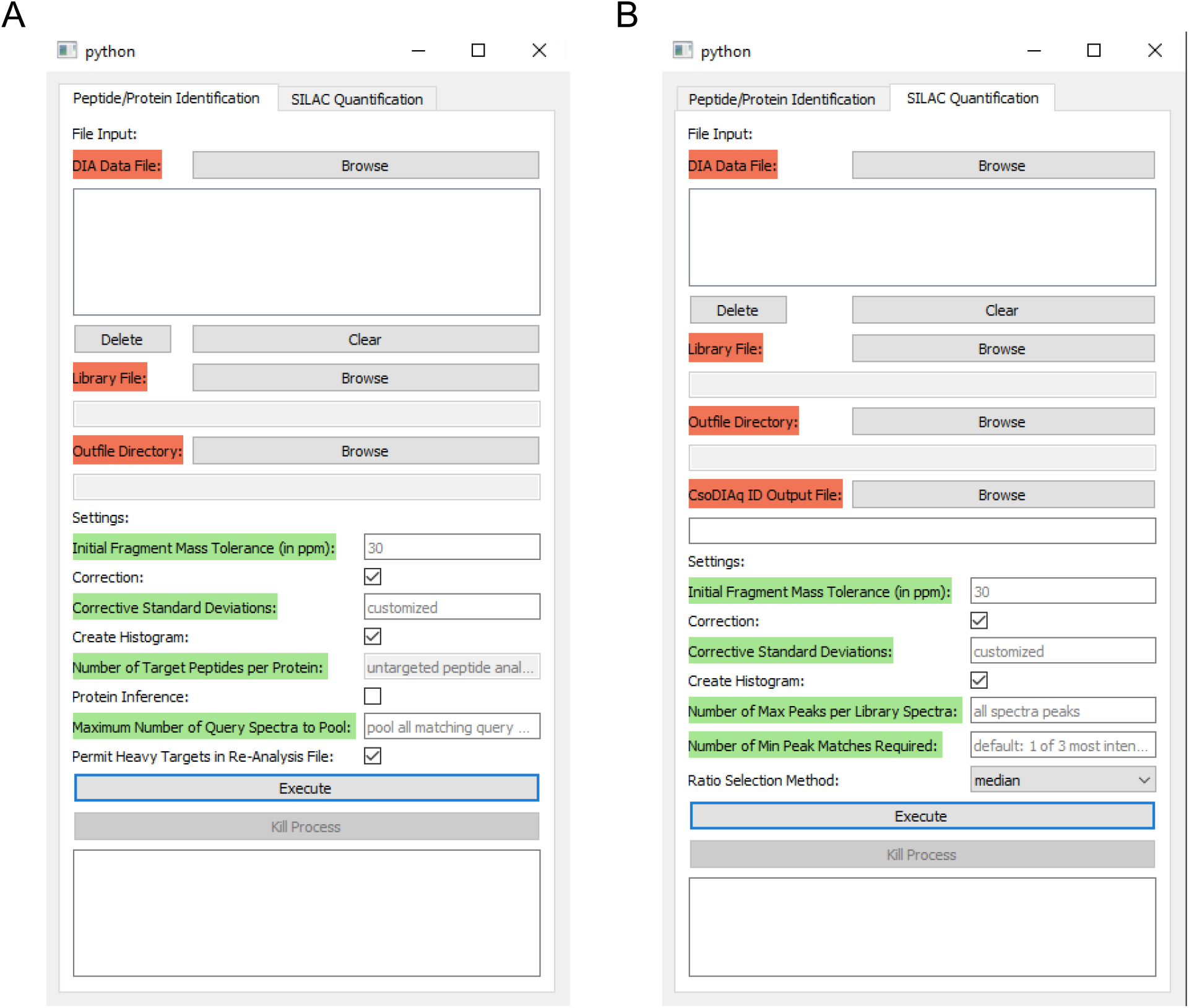
Graphical User Interface layout. **A**, GUI settings for peptide and protein identification step. **B**, GUI settings for SILAC quantification step.

## DISCUSSION

The CsoDIAq software package introduced here enables the first unified solution to DISPA data analysis, which we expect to enable more widespread adoption. Furthermore, the applicability of CsoDIAq to standard LC-MS DIA analyses expands the utility. CsoDIAq introduces several algorithmic advances. These include spectra pooling, which significantly reduces the time required to complete a DIA-based analysis. Further, the MaCC score and PPM correction concepts can apply to similar tools in future development. By combining these techniques with the projected spectrum scoring concept outlined in the MSPLIT-DIA package, we demonstrated an overall enhancement in the quantity of peptides and proteins identified. Further, the results demonstrate that the CsoDIAq can accurately quantify peptides and proteins. Finally, CsoDIAq greatly increases usability through the addition of a GUI. Altogether, the CsoDIAq software package simplifies and enhances DISPA data analysis.

## ASSOCIATED CONTENT

### Supporting Information

The Supporting Information is available free of charge

Supplementary figures, DOCX

Supplementary Table 1, XLSX

## Author Contributions

Conceptualization, J.G.M and C.W.C.; Software, C.W.C; Validation, C.W.C; Formal Analysis, C.W.C; Investigation, J.G.M; Resources, J.G.M.; Data Curation, C.W.C; Writing – Original Draft, C.W.C; Writing – Review & Editing, J.G.M and C.W.C.; Visualization, C.W.C; Supervision, J.G.M.

## ACKNOWLEDGMENT

We thank David Tabb for guidance in understanding the IDpicker algorithm. We thank Dasom Hwang for graphic design assistance.

