## supplemental figures for "CsoDIAq Software for Direct Infusion Shotgun Proteome Analysis (DISPA)"

**Table of Contents:**

**Figure S1.** Original Data Analysis Workflow Flowchart

**Figure S2.** Fragment ion intensities measured for isolation widths 0.4, 1.0, and 2.0 m/z for the peptides AIEIVDQALDR and ETDIGVTGGGQGK

**Table S1.** Comparison of peptide and protein identifications by either CsoDIAq or MSPLIT-DIA using DISPA or LC-MS data matching figure 4.

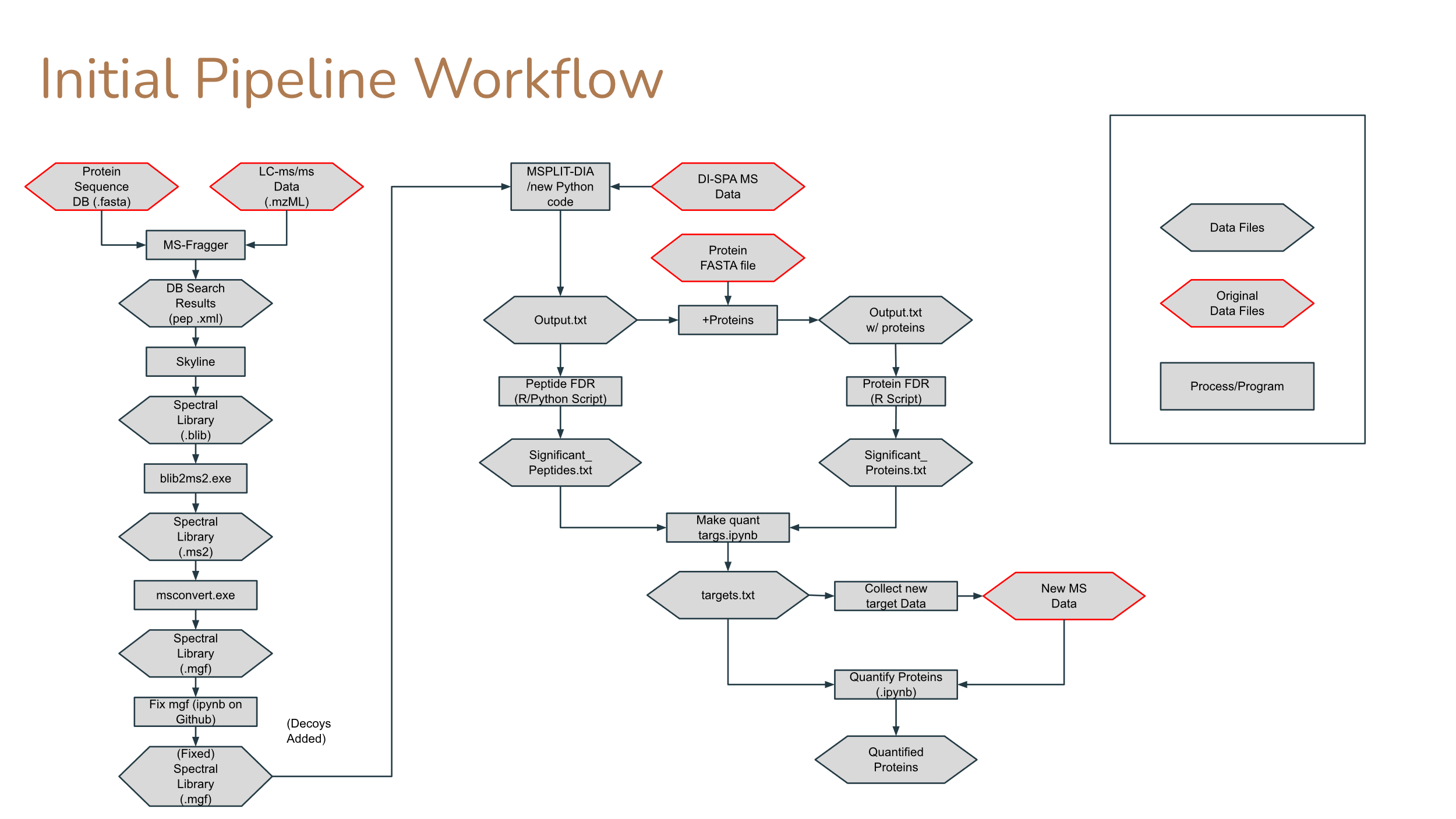

**Figure S1: Original Data Analysis workflow (from Meyer *et al.*, Nature Methods, 2020).** This workflow diagram indicates the complexity of analysis relative to the new analysis enabled by CsoDIAq shown in Figure 1.

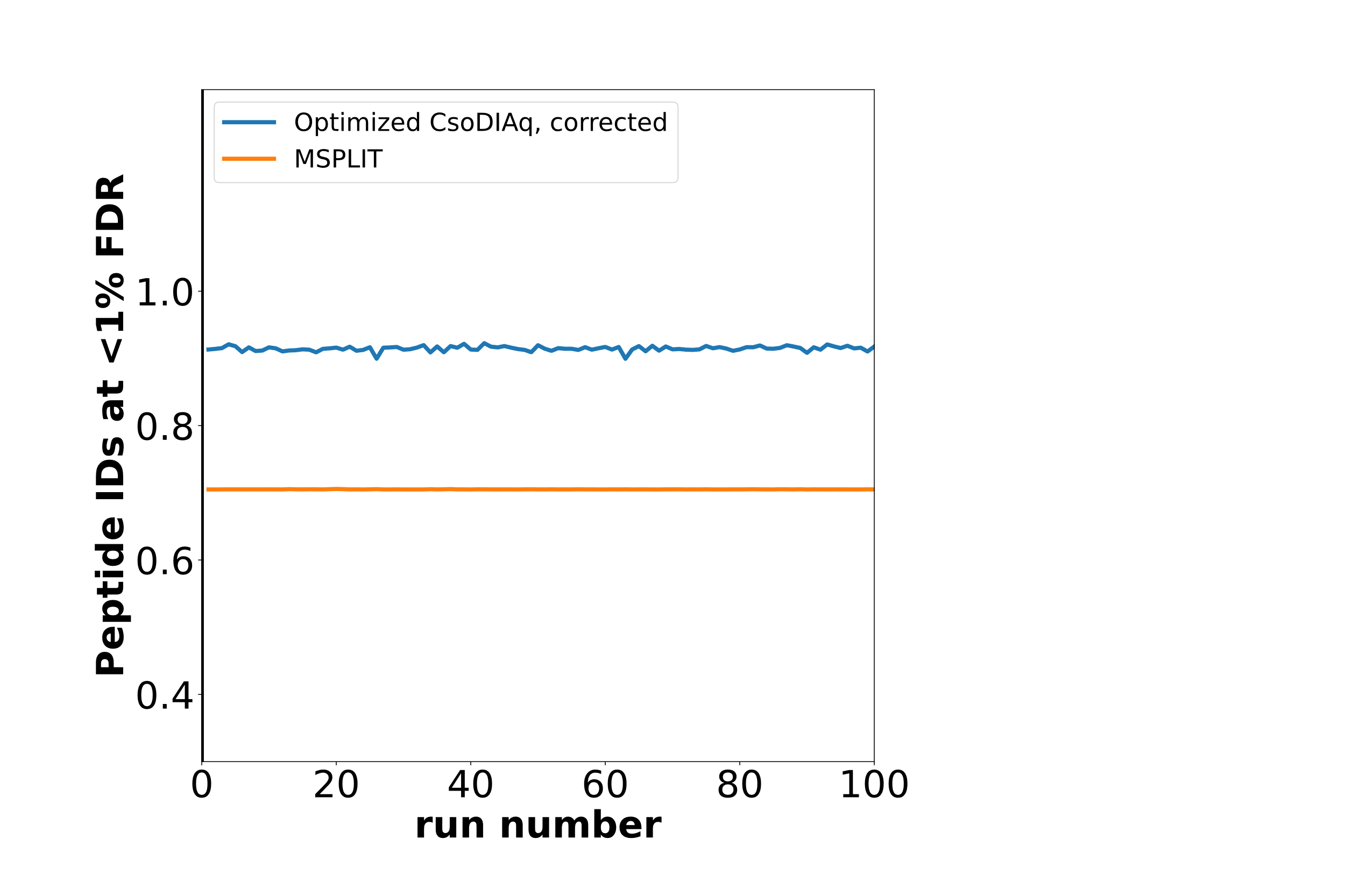

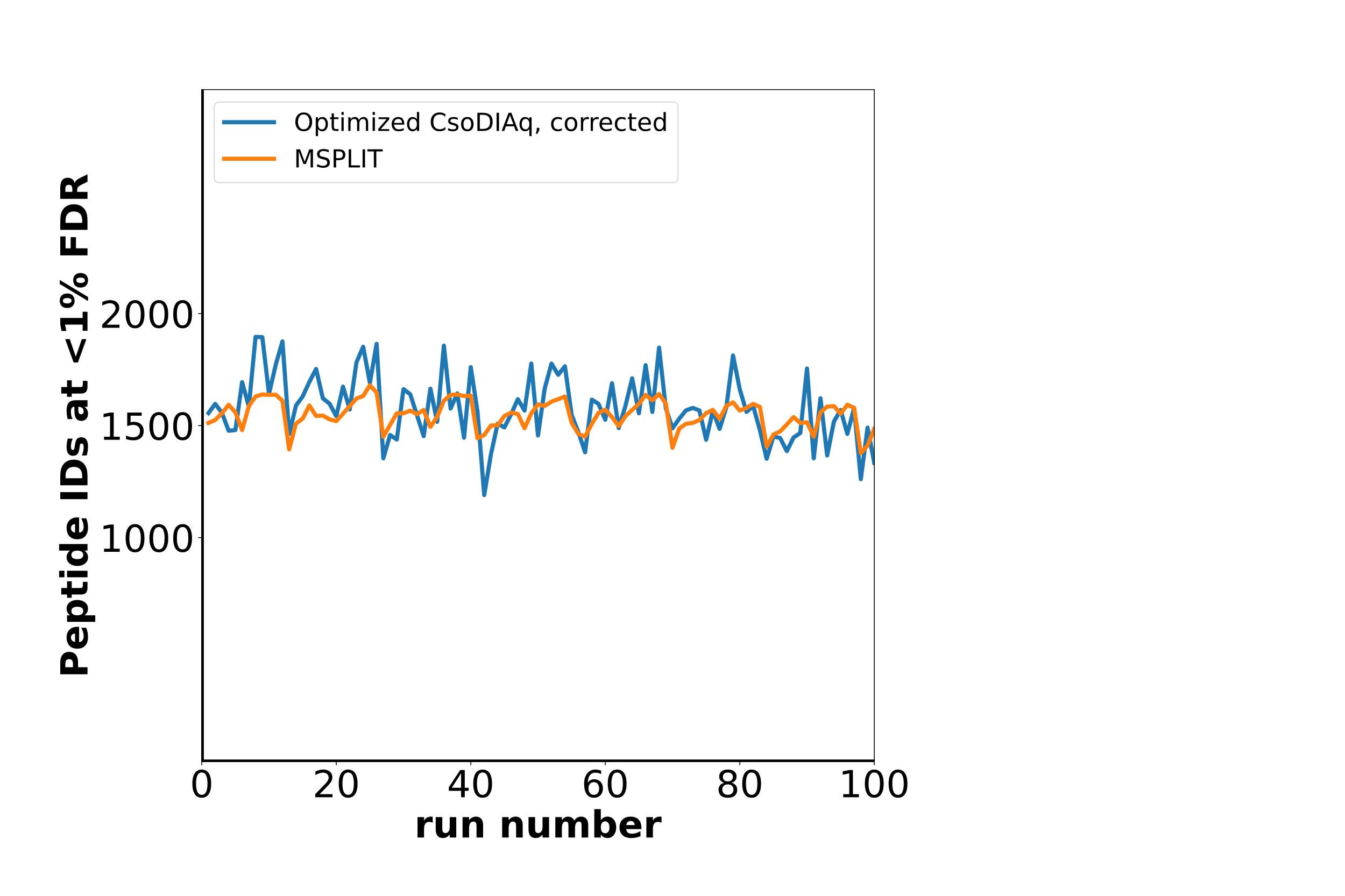

B

A

**Figure 2: Direct comparison of MSPLIT and CsoDIAq using the same MGF library.** (A) Number of peptide identifications across the 100 replicate DISPA replicates dataset. (B) Minimum cosine score for each replicate across the 100 replicate DISPA replicates dataset.
